# Hydroxymethylated-*P16* Allele Is Transcription-Inactive

**DOI:** 10.1101/405522

**Authors:** Ying Gan, Paiyun Li, Xiao Han, Sisi Qin, Chenghua Cui, Zhaojun Liu, Jing Zhou, Liankun Gu, Zhe-ming Lu, Baozhen Zhang, Dajun Deng

**Affiliations:** Key Laboratory of Carcinogenesis and Translational Research (Ministry of Education/ Beijing), Division of Etiology, Peking University Cancer Hospital and Institute, Fu-Cheng-Lu #52, Haidian District, Beijing, 100142, China

**Keywords:** *P16* gene, CpG islands, methylation, hydroxymethylation, epigenetic editing

## Abstract

**Background:** 5-Methylcytosine can be oxidized into 5-hydroxymethylcytosine (5hmC) in the genome. Methylated-*P16* (P16M) can be oxidized into completely hydroxymethylated-*P16* (P16H) in human cancer and precancer cells. The aim of this study is to investigate the biological function of P16H.

**Methods:** True P16M and P16H were analyzed using bisulfite/TAB-based assays. A ZFP-based *P16-*specific dioxygenase (P16-TET) was constructed and used to induce P16H. Cell proliferation and migration were determined with a series of biological analyses.

**Results:** (**A**) The 5hmCs were enriched in the antisense-strand of the *P16* exon-1 in HCT116 and AGS cells containing methylated*-P16* alleles (P16M). (**B**) P16-TET induced both P16H and *P16* demethylation in H1299 and AGS cells and reactivated *P16* expression. Notably, P16H was only detectable in the sorted P16-TET H1299 and AGS cells that did not show *P16* expression. (**C**) P16-TET significantly inhibited the xenograft growth derived from H1299 cells in NOD-SCID mice, but did not inhibit the growth of *P16-*deleted A549 control cells. *P16-*siRNA knockdown could rescue P16-TET-inhibited cell migration.

**Conclusion:** Hydroxymethylated *P16* alleles are transcriptionally inactive.

**AUTHOR SUMMARY:** It is well known that 5-methylcytosine (5mC) in genomic DNA of mammalian cells can be oxidized into 5-hydroxymethylcytosine (5hmC) and other derivates by DNA dioxygenase TETs. While conversion of 5mC to 5hmC plays an important role in active DNA demethylation through further oxidations, a certain proportion of 5hmCs remain in the genome. Although it is supposed that occurrence of 5hmCs may contribute to the flexibility of chromatin and the protection of the bivalent promoters from hypermethylation, the direct effect of 5hmCs on gene transcription is unknown. In the present study, we engineered a zinc-finger protein-based P16-specific DNA dioxygenase and used it to induce *P16* hydroxymethylation and demethylation in cancer cells. Our results demonstrate, for the first time, that the hydroxymethylated *P16* alleles retain transcriptionally inactive. This is supported by our recent findings that mRNAs are always transcribed only from the unmethylated P16 strands, but not from the hydroxymethylated/methylated strands in HCT116 cells, and that the risks for malignant transformation are similar for patients with the *P16* methylation-positive oral epithelial dysplasia with and without *P16* hydroxymethylation in a prospective study.

## INTRODUCTION

It is well known that ten-eleven translocation methylcytosine dioxygenases (TET-1/2/3) oxidize 5-methylcytosine (5mC) to 5-hydroxymethylcytosine (5hmC), 5-formylcytosine (5fC), and 5-carboxylcytosine (5caC) in the genome [1–4]. While oxidation of 5mC leads to active DNA demethylation, a certain proportion of 5hmC sites remain in the genome with a strand-asymmetric and strand-symmetric distribution pattern that provides its own regulatory function [5–9]. Although it is frequently reported that the 5hmC level of some genes is positively correlated with increased gene expression, it is not clear whether 5hmC itself or related DNA demethylation contribute to the reactivation of gene transcription.

Typical bisulfite-based assays cannot discriminate 5mCs from 5hmCs. The classic term “DNA methylation” is, in fact, total DNA methylation, including true methylation and hydroxymethylation. Total methylation of the CpG island (CGI) flanking the transcription start site (TSS) in the *P16* gene (*CDKN2A*) is prevalent in human cancer and precancerous tissues [10,11] and is linked to increased cancer development from epithelial dysplasia in many organs [12–18]. *P16* methylation (P16M) not only directly inactivates *P16* transcription [19] but also represses *ANRIL* transcription [20]. Our recent study demonstrated that there were dense 5hmCs in the *P16* exon-1 CGI in HCT116 cells, and no mRNA transcripts from the hydroxymethylated *P16* (P16H) alleles were detected in the cells [21,22]. P16H was detected in 9.3% of human oral epithelial dysplasia (OED) tissues [23]. However, the malignant transformation risk was similar between P16M-positive OED patients with and without P16H. It is a fundamental question in epigenetic research to clarify whether hydroxymethylation of TSS-flanking CGIs leads to transcriptional activation of genes.

In this study, we characterized the distribution patterns of 5hmCs within the sense and antisense strands (S- and AS-strands) of the *P16* promoter and exon-1 CGIs using detailed TET-assisted bisulfite (TAB)-based assays, and found that 5hmCs were enriched in the AS-strands of *P16* exon-1 CGIs in cancer cells. To elucidate the possible role of P16H, an epigenetic editing tool, *P16*-specific TET-1 (P16-TET), was constructed and used to induce P16H in cancer cell lines. Notably, our data showed, for the first time, that P16H itself could not reactivate gene transcription.

## RESULTS

### Characterization of 5hmCs in the *P16* Exon-1 CGI

We recently found that there were dense 5hmCs in the *P16* exon-1 in HCT116 cells [21], in which the wildtype *P16* alleles are silenced by DNA methylation and the mutant alleles containing a G-insertion in exon-1 are unmethylated. To characterize the distribution pattern of 5hmCs in the *P16* CGI, the S- and AS-strands of the *P16* promoter and the exon-1 regions were amplified using conventional bisulfite-modified and TAB-modified single-strand DNA samples from HCT116 cells as templates. Next, the proportions of total P16M- and P16H-containing fragments were quantitatively analyzed by denaturing high performance liquid chromatography (DHPLC). As expected, the total P16M peak and the P16U peak were both detected in the all bisulfite PCR products from both the S- and AS-strands of the *P16* promoter and exon-1 fragments (Figure 1A-D: HCT116, left charts). However, a high P16H peak was detected only in the exon-1 AS-strand and the P16H proportion reached up to 88% (=0.77/0.87) (Figure 1D: HCT116_TAB, left chart). In the promoter AS-strand fragment, the P16H peak was very low (Figure 1C: HCT116_TAB, left chart). In the S-strands of the promoter and exon-1 fragments, much lower levels of the P16H peaks were detected (Figures 1A and 1B: HCT116_TAB, left charts).

**Figure 1.**
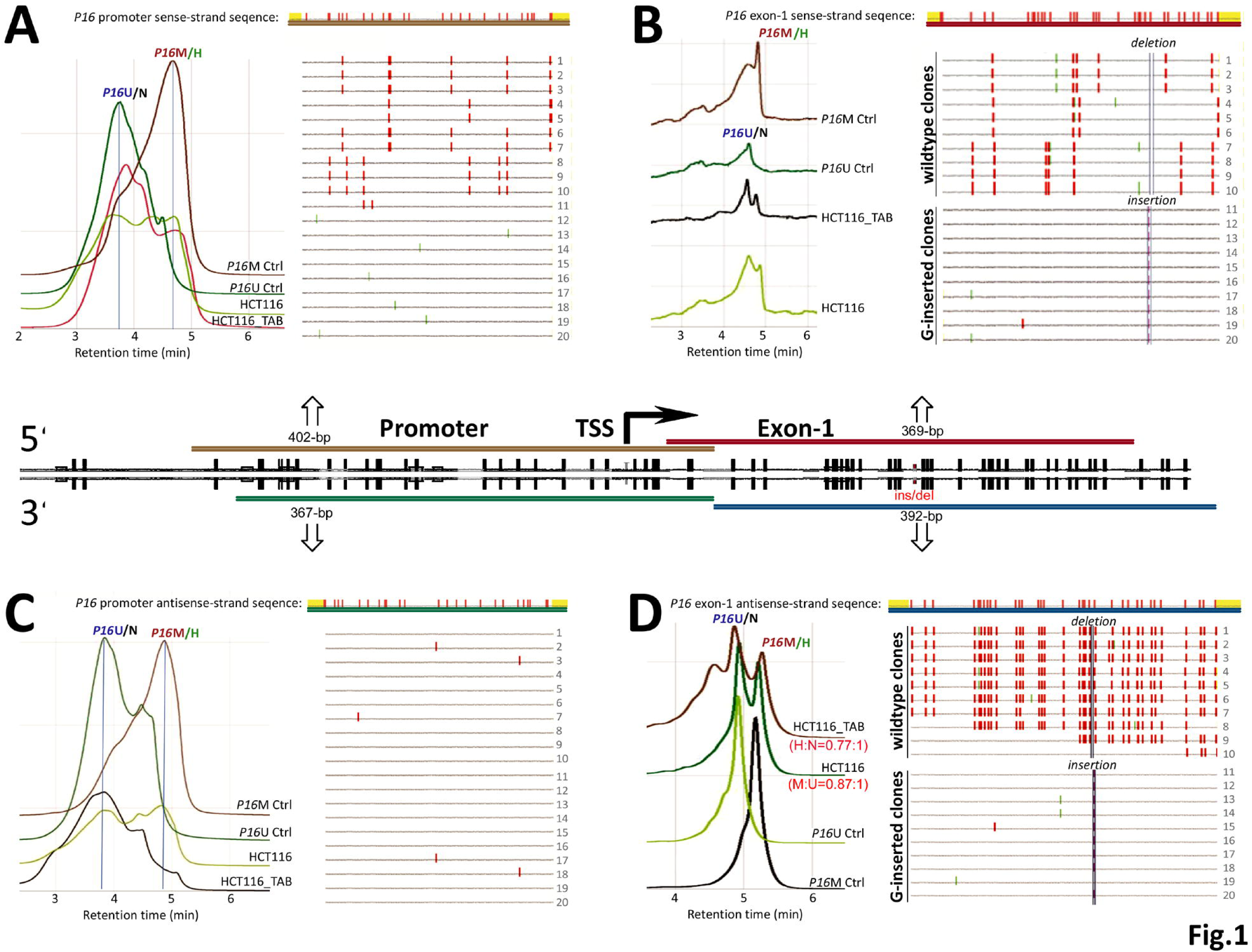
TAB-DHPLC and TAB sequencing detected 5hmCs in the sense- and antisense-strands of *P16* promoter and exon-1 of HCT116 cells. (***A****-**D***) Results of four amplicons in the sense- and antisense-strands of the *P16* promoter and exon-1 regions are illustrated in the middle chart; Left charts: chromatograms of bisulfite- and TAB-PCR products for four strands. P16M Ctrl and P16U Ctrl: the corresponding PCR products of *P16-*methylated RKO cells and *P16-*unmethylated MGC803 cells; Right images: Results of clone sequencing of HCT116_TAB PCR products for four strands. 5hmC density: 82.9% (290/350) and 17.6% (58/330) for wildtype exon-1 antisense- and sense-strand clones, and 2.0% (5/250) and 22.3% (58/260) for promoter antisense- and sense-strand clones. Each line represents one clone, respectively. Each red dot represents one 5hmC. Green dots indicate TAB-unmodified cytosines. The locations of G-insertion and G-deletion in exon-1 are also labeled. The amplicon sequences of four strands are placed at the top of the images. M:U and H:N, ratios of the peak height for <u>M</u>ethylated-*P16* to <u>U</u>nmethylated-*P16* alleles and <u>H</u>ydroxymethylated-*P16* to <u>N</u>ot hydroxymethylated-*P16* alleles in bisulfite-DHPLC and TAB-DHPLC analyses, respectively.

The TAB sequencing results for the HCT116_TAB PCR products confirmed the DHPLC analysis results (Figure 1A-D: right charts). Dense 5hmCs were found in the wildtype exon-1 AS-strand (tracked with a G-deletion; 5hmC-density, 82.9%), but not in the paired S-strand (5hmC-density, 17.6%). No clone containing more than one 5hmC was detected in the promoter AS-strand (5hmC density, 2.0%). Sporadic 5hmCs were distributed in the promoter S-strand (5hmC density, 22.3%). Together, the results of the TAB-DHPLC and TAB sequencing analyses consistently demonstrated that 5hmCs were enriched mainly in the AS-strand of the wild-type *P16* exon-1 in HCT116 cells. This indicates that wild-type *P16* exon-1 is hydroxymethylated mainly in the AS-strand and is methylated truly in the S-strand in HCT116 cells. As described below, dense 5hmC sites were also detected in the AS-strand of the *P16* exon-1 in gastric cancer AGS cells.

### Construction of Engineered P16-TET

To study whether P16H affects gene transcription, an expression controllable *P16-*specific dioxygenase P16-TET and its inactive mutant control vector were constructed through fusing an engineered *P16* promoter-specific seven zinc finger protein (7ZFP-6I) [24] with the catalytic domain of human TET1 and integrated into the pcDNA3.1 expression vector (Figure S2A). H1299 cells were chosen because epigenetic editing of the methylated *P16* CGIs by the *P16*-specific transcription factor (P16-ATF; 7ZFP-6I-VP64) has been optimized in this cell type [24]. As expected, the results of both qRT-PCR and immunofluorescence staining showed that the methylated *P16* alleles were re-activated in H1299 cells 6 days after transient transfection with the P16-TET vector (Figure 2). Such *P16* reactivation was not observed in the P16-TET mutant control cells. This indicates that P16-TET is *P16* gene reactive and could be used in further studies.

**Figure 2.**
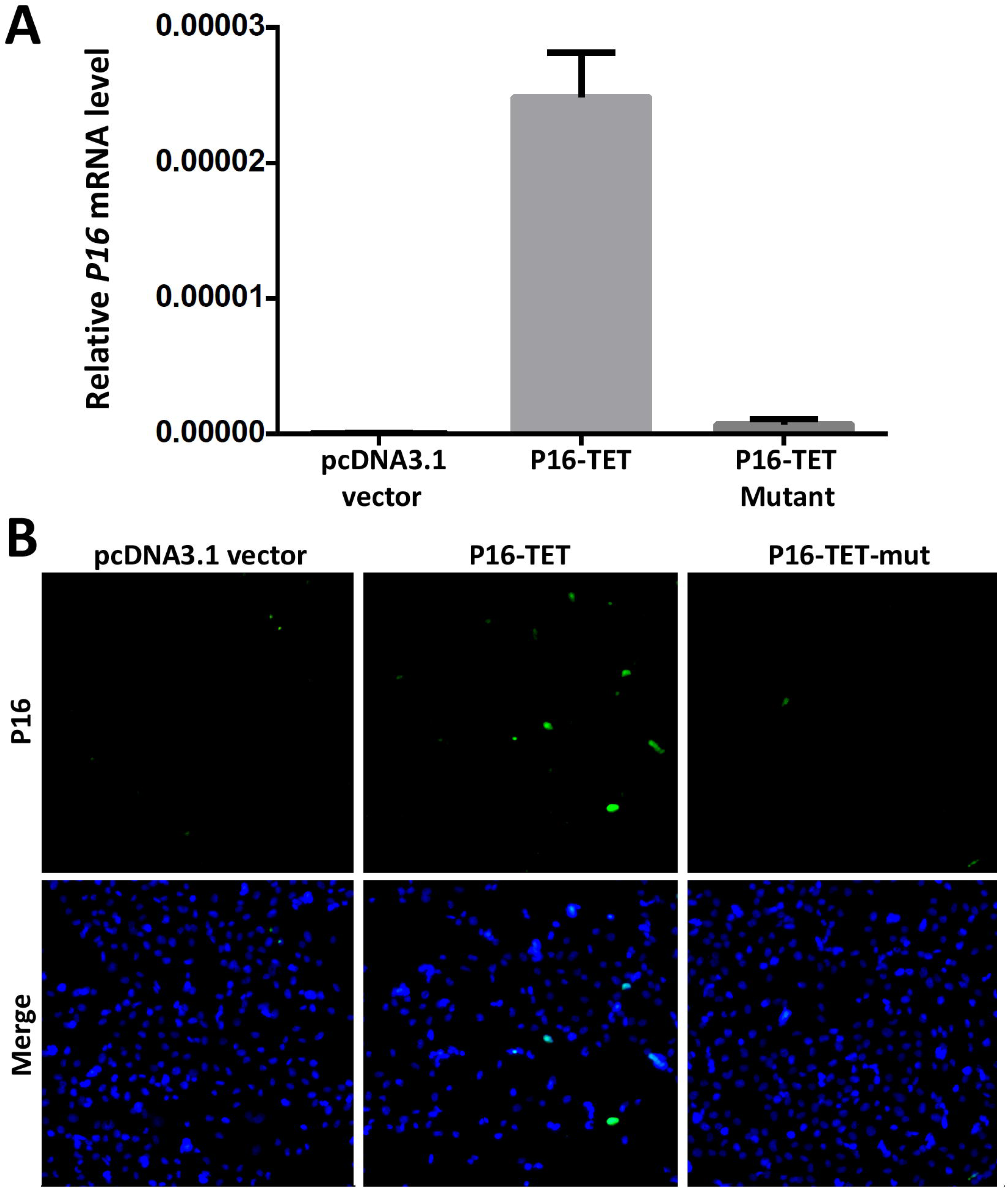
Reactivation of methylated *P16* alleles in H1299 cells 6 days after P16-TET transient transfection. (**A**) qRT-PCR. (**B**) Immunofluorescence staining.

### Induction of P16H by P16-TET

To study the possible biological functions of *P16*-specific hydroxymethylation, the P16-TET coding sequence was further integrated into the pTRIPZ lentivirus vector carrying a “Tet-on” switch to allow the gene expression to be controlled for stable transfection (Figure S2B). In the P16-TET stably transfected H1299 cells, the results of the TAB methylation-specific PCR (MSP) analysis showed that P16H signals appeared in the P16-TET cells 3 days after the induction of doxycycline (Dox; final conc. 0.25 μg/mL) (P16-TET&Dox_3d; Figure 2A, TAB-MSP), but did not appear in cells transfected with the empty vector (control cells with Dox treatment) (Vector&Dox_14d) or in baseline P16-TET cells without Dox induction, in which only nonhydroxymethylated *P16* alleles (P16N) were detected. In the MSP analysis, P16U was detectable in the P16-TET&Dox cells 3 days following Dox induction (Figure 3A, MSP). The bisulfite-DHPLC results showed that a low P16U peak was detected beginning on the 14^th^ day (Figure S3A, red-arrow). Two P16U clones were also observed on the 28^th^ day from the bisulfite sequencing (Figure S3B, red-star). These results indicate that both P16H and P16U were induced in the P16-TET&Dox cells.

**Figure 3.**
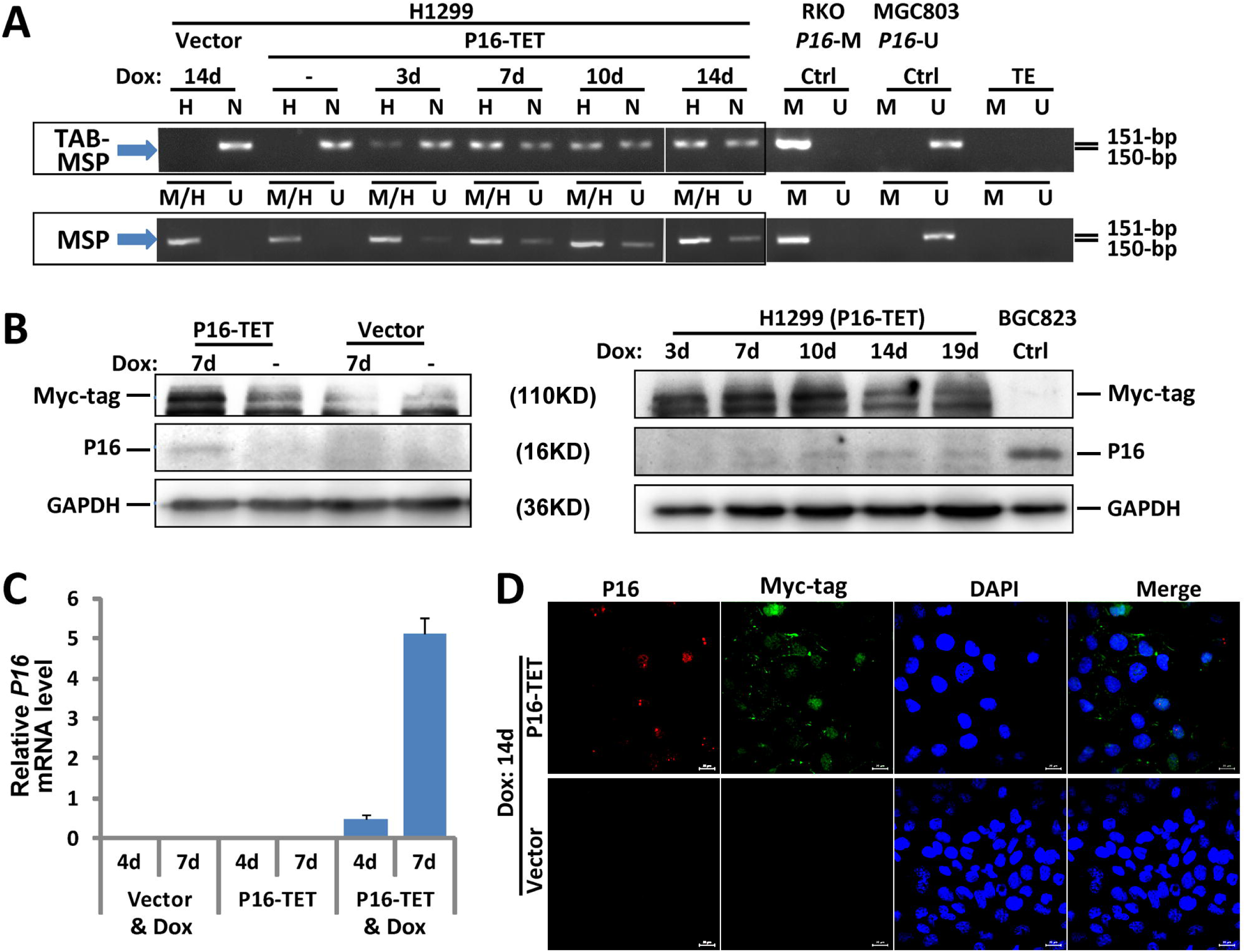
P16-TET induces hydroxymethylation of *P16* CpG islands and reactivates expression of methylated *P16* alleles in H1299 cells. (**A**) TAB-MSP analysis for detecting hydroxymethylated (H)- and nonhydroxymethylated (N)-*P16* CpG alleles in H1299 cells stably transfected with P16-TET or empty control vector after doxycycline treatment. The MSP analysis results were also listed. Genomic DNA from RKO and BGC832 cells was used as P16M and P16U controls in the MSP assays, respectively. (**B**) Western blot analysis for detecting the P16 protein; Dox (+/-): with or without the doxycycline treatment (final conc. 0.25 μg/mL). Proteins from BGC832 cells were used as a P16U/active control. (**C**) qRT-PCR results for detecting *P16* mRNA levels relative to *Alu* RNA levels; (**D**) Immunofluorescence confocal analysis for detecting P16 expression.

Furthermore, the Western blot results revealed that P16 protein was detected in the P16-TET&Dox cells since the 7^th^ day, but not in the Vector&Dox control cells (7d; Figure 3B). The qRT-PCR results showed a weak reactivation of *P16* transcription beginning on the 4^th^ day (Figure 3C). The immunofluorescence confocal microscopy results confirmed the presence of P16 protein in the nuclei of H1299 cells (Figure 3D). In addition, the expression status of the control genes *P15* and *P14* was not affected, whereas the expression level of *ANRIL,* which is coordinately expressed with *P16*, was increased (Figure S4). This suggests a high specificity for the zinc finger protein-based P16-TET to induce P16H and P16U.

Similarly, on the 7^th^ day after Dox induction, transcriptional reactivation of *P16* was also observed in P16-TET stably transfected gastric cancer AGS cells, in which *P16* alleles are homogenously methylated (Figures 4A-4E). Interestingly, P16U signals were not detected in P16-TET AGS cells after Dox induction for 11 days (P16-TET&Dox_11d) in the bisulfite-DHPLC and bisulfite sequencing analyses (Figure 4A and 4C). P16H signals were observed in the TAB-DHPLC and TAB sequencing results (Figure 4B and 4D), indicating that hydroxymethylation occurred earlier than demethylation at P16 CGIs. A few baseline 5hmCs were also found in the *P16* exon-1 AS-strand of AGS mock control cells. Although weak *P16* mRNA signals were detected in P16-TET AGS cells after Dox induction for 7 days and 11 days according to sensitive RT-PCR analysis (Figure 4E), P16 protein was not detected in these cells according to the insensitive Western blot analysis (Figure 4F).

**Figure 4.**
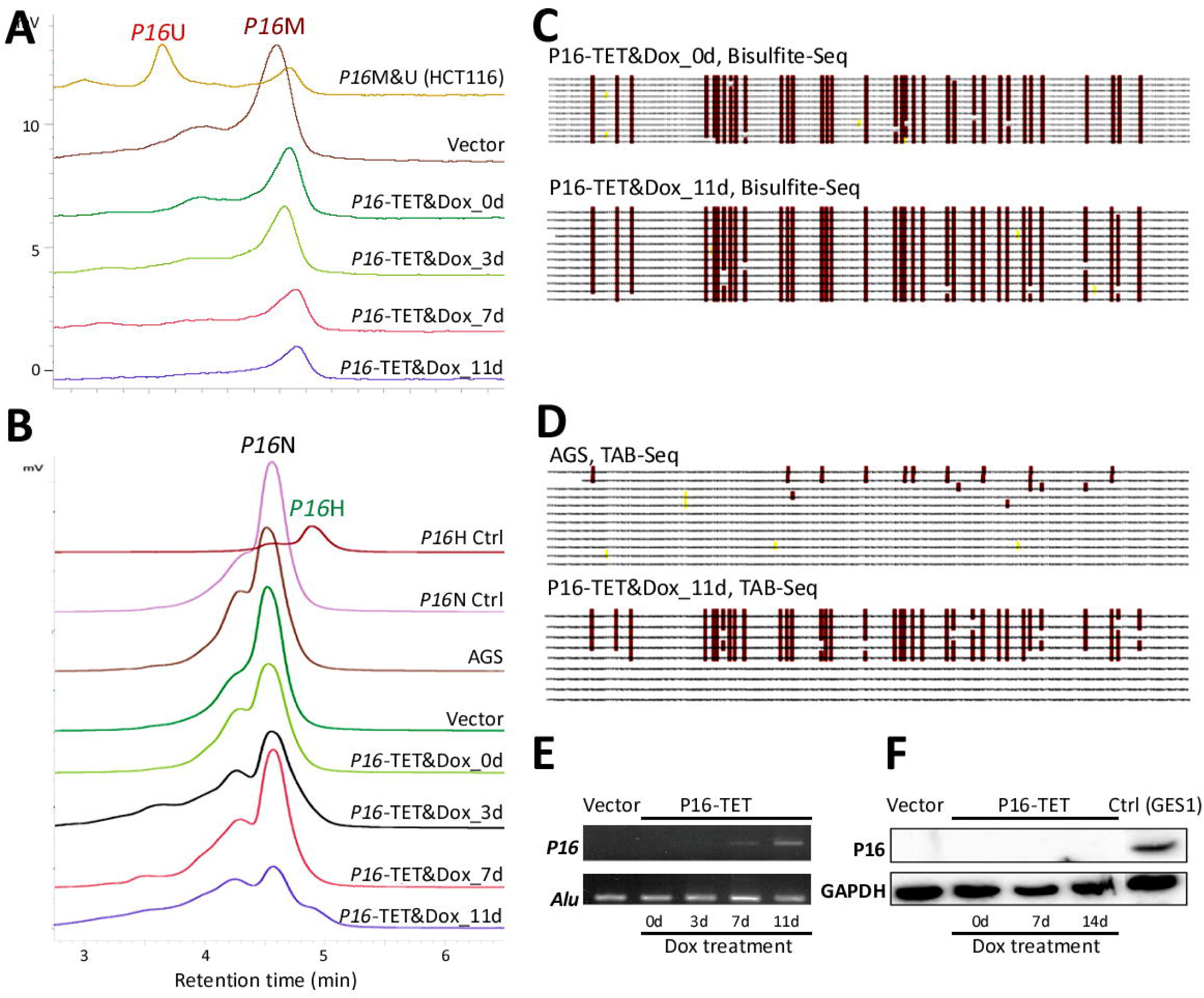
P16-TET induces hydroxymethylation of *P16* CpG islands and reactivates expression of methylated-*P16* alleles in AGS cells. (**A**) Bisulfite-DHPLC analysis for detecting methylated-*P16* (P16M) and unmethylated-*P16* (P16U) PCR products for the exon-1 antisense strand in P16-TET-transfected AGS cells with different doxycycline induction times. (**B**) The TAB-DHPLC analysis detected the hydroxymethylated *P16* (P16H) PCR products and nonhydroxymethylated *P16* (P16N) PCR products. (**C** and **D**) Bisulfite and TAB sequencing for detecting 5mC and 5hmC sites, respectively, in the same PCR products as were analyzed by DHPLC. (**E** and **F**) The results of RT-PCR and Western blot analysis for detecting P16 reactivation in AGS cells.

### Transcription Silencing of *P16* alleles by Hydroxymethylation

To clarify whether DNA hydroxymethylation or demethylation contributes to *P16* reactivation, we further analyzed the hydroxymethylation status of *P16* CGIs in cell subpopulations with strong, weak, and no P16 staining (P16(+), P16(±), and P16(-)) that were sorted from P16-TET&Dox_21d H1299 cells (Figure 5A). Interestingly, P16H signal was detected only in the P16(-) subpopulation, but not in the P16(+) and P16(±) subpopulations in the TAB-MSP analysis (Figure 5B). TAB sequencing also showed dense 5hmCs among 3 of the 14 clones (21.4%) of the exon-1 AS-strand TAB-PCR products from the P16(-) subpopulation, with an average hydroxymethylation density of 95.2% for these 3 clones (Figure 5C). The occurrence of 5hmCs in the promoter AS-strand was not detected in the TAB-DHPLC and TAB sequencing results (data not shown).

**Figure 5.**
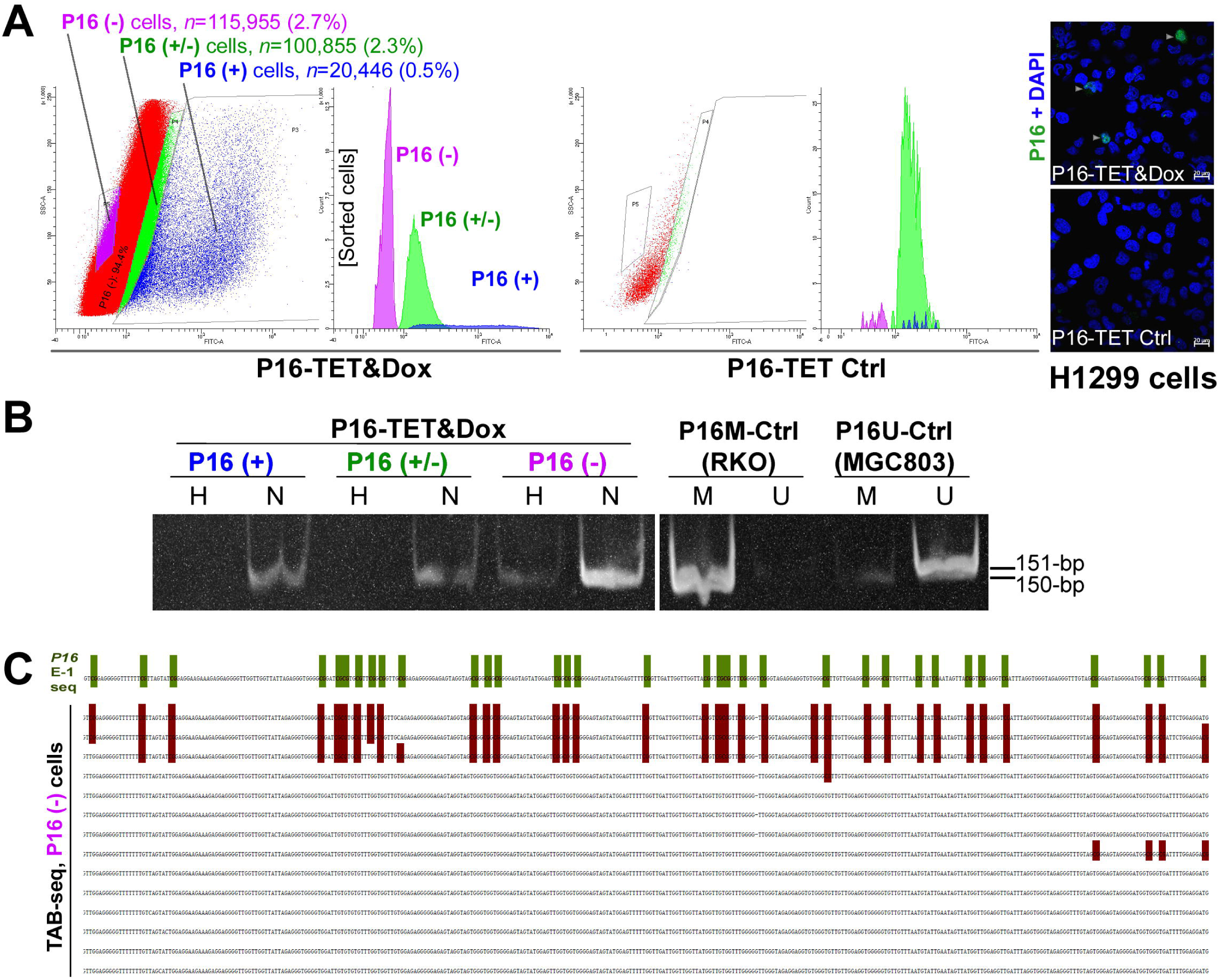
Characterization of P16H in FACS-sorted subpopulations of H1299 cells with various levels of of *P16* expression reactivation. (**A**) FACS sorting of P16-TET stably transfected H1299 cells with and without Dox treatment. The confocal images of the P16 protein staining status are also attached. (**B**) Detection of the DNA hydroxymethylation status of *P16* alleles in various FACS-sorted H1299 subpopulations with strong, weak, and no P16 immunostaining (P16(+)/(±)/(-)) in the TAB-MSP analysis. (**C**) The results of TAB sequencing for the *P16* CpG islands in the P16-negative subpopulation.

The above results were further confirmed in AGS cells. As described above, P16 protein could not be detected in P16-TET&Dox AGS cells after Dox treatment for 11 days (Figure 4F). To obtain a P16(+) AGS subpopulation by FACS, the DNA methyltransferase inhibitor 5-aza-deoxycytidine (DAC, final concentration 20 nmol/L) was used to increase the P16 protein level within P16-TET AGS cells. In the immunostaining cell analysis, nucleic P16 protein was detected in 3.5% of P16-TET AGS cells after DAC treatment for 10 days (P16-TET&DAC_10d, with baseline P16-TET expression without Dox induction), while nucleic P16 protein was detected in only 0.5% of the AGS cells treated with DAC alone (Figure S5). Next, the P16(+), P16(±), and P16(-) subpopulations were sorted from these P16-TET&DAC_10d AGS cells (Figure 6A). Once again, the P16H signal was detected only in the P16(-) subpopulation, and not in the P16(+) and P16(±) cells by the TAB-MSP and TAB-DHPLC assays (Figures 6B and 6C). In contrast, P16N signal was detected in all three subpopulations. The TAB sequencing results confirmed this. Dense 5hmCs were observed in the *P16* exon-1 AS-strand in the P16(-) subpopulation, but not in the P16(+) subpopulation (Figure 6E).

**Figure 6.**
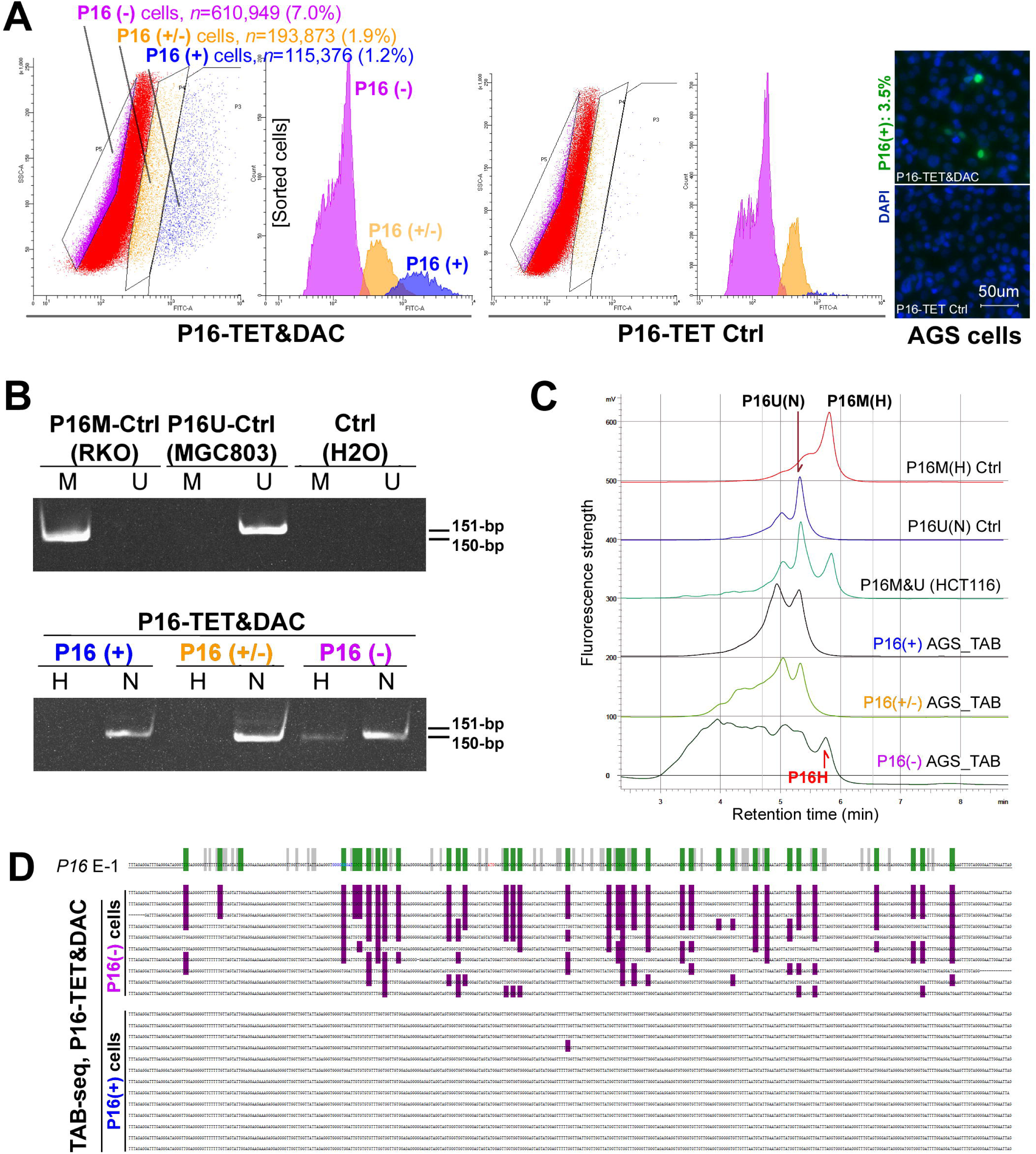
Characterization of P16H in FACS-sorted subpopulations of AGS cells with various levels of *P16* expression reactivation. (**A**) FACS-sorting of P16-TET stably transfected AGS cells with and without DAC treatment. The confocal images of the P16 protein staining status are also attached. (**B**) Detection of the DNA hydroxymethylation status of *P16* alleles in various FACS sorted AGS subpopulations with strong, weak, and no P16-immunostaining (P16(+)/(±)/(-)) in the TAB-MSP analysis. (**C**) The results of TAB-DHPLC for the *P16* CpG islands in three subpopulations. (**D**) The results of TAB sequencing for the antisense strand of *P16* exon-1 in P16(+) and P16(-) subpopulations.

Collectively, the above results indicate that P16H occurs only in P16(-) cells, and not in P16(+) and P16(±) cells, suggesting that the P16H alleles should be transcriptionally inactive.

### P16 Allele-Dependent Inhibition of Tumor Growth by P16-TET

Although a proliferation difference was not observed between the P16-TET and control vector, which were stably transfected in H1299 cells *in vitro* (Figure 7A), the average weight of tumor xenograftsof the P16-TET stably transfected cells was significantly lower than that of the control cells in NOD-SCID mice (*n*=8) on the 50^th^ post-transplantation day (*P*<0.001, Figures 7B and 7C). Morphologic differences were not observed between P16-TET and control vector xenografts (Figure 7D). This result was confirmed in a repeat experiment (Figure S6A). Meanwhile, this difference could not be observed in xenograft tumors from lung cancer A549 control cells in which the *P16-P15-P14* alleles were homogeneously deleted (Figure S6B). These data suggest that P16-TET may specifically inhibit the growth of cancer cells *in vivo* in a *P16* allele-dependent manner.

**Figure 7.**
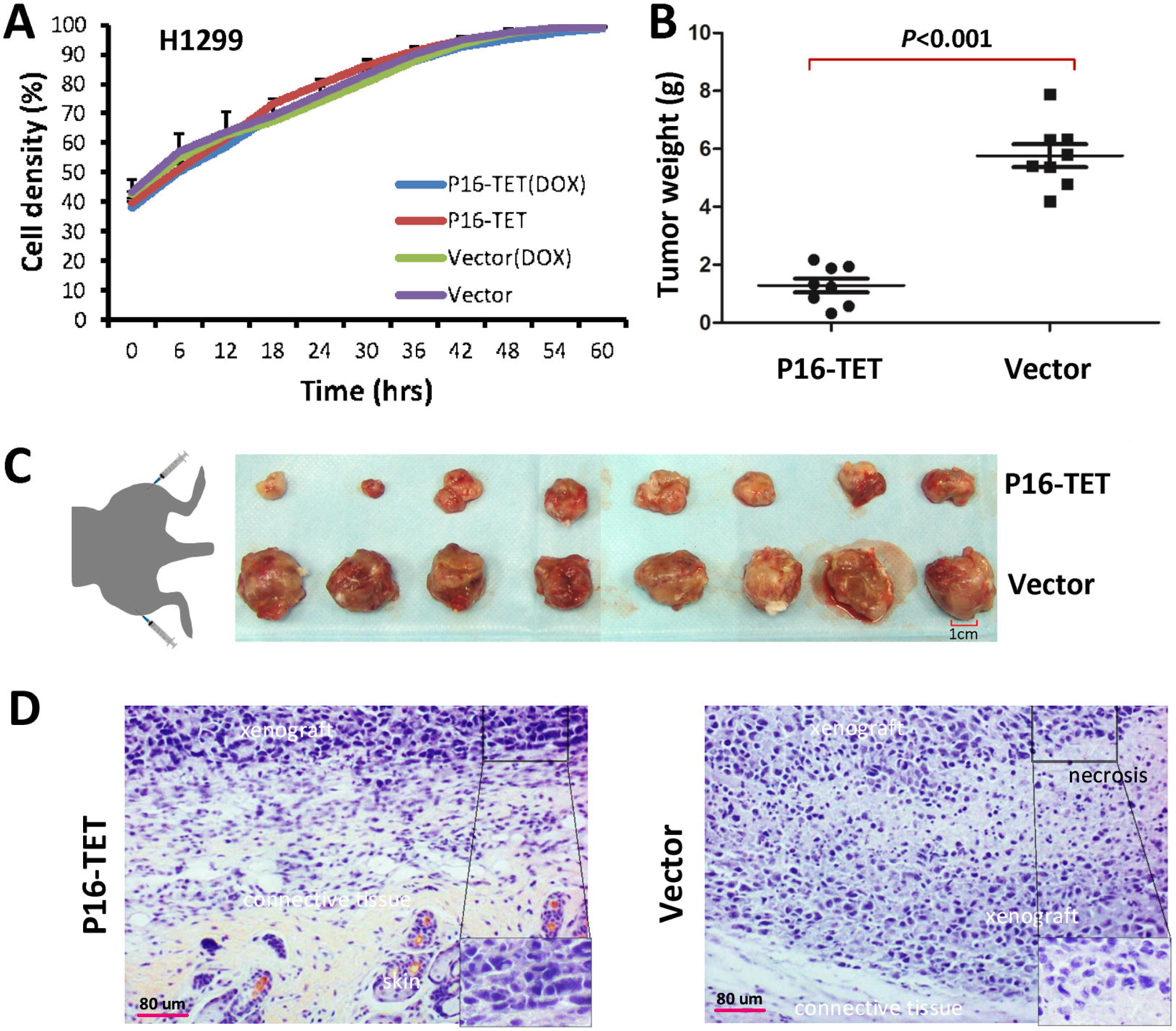
Effects of P16H on the proliferation of H1299 cells *in vitro* and *in vivo*. (***A***) Cell proliferation curves for H1299 cells with and without P16-TET expression in a live content kinetic imaging platform; (**B**) Comparison of weights of H1299 tumor xenografts with and without stable P16-TET transfection in SCID mice; (**C** and **D**) Images of xenografts on the 50^th^ experimental day.

Although P16-TET did not affect the proliferation of H1299 cells *in vitro*, the results of the IncuCyte ZOOM wound-scratch and typical transwell assays showed that P16-TET significantly inhibited H1299 cell migration (Figures S7A and S7B). In a rescue assay, *P16* siRNA-knockdown significantly reversed the inhibited migration of the P16-TET&Dox H1299 cells (Figure S7C). These results provide further evidence to support that P16-TET may inhibit cell migration through *P16* reactivation.

## DISCUSSION

DNA hydroxymethylomes at the base-resolution level have been analyzed in embryonic stem cells, adult tissues, and tumors [25–31]. Many functions of DNA hydroxymethylation in the genome have been illustrated by *TET-1/2/3* knockout studies [5–7,27,31,32]. However, the actual effect of hydroxymethylation of CGIs on gene transcription remains elusive. In the present study, we demonstrated that 5hmCs were enriched in the AS-strand of the *P16* exon-1 CGI. Most importantly, this study showed for the first time that DNA hydroxymethylation itself could not reactivate *P16* gene transcription. Instead, hydroxymethylation-mediated active DNA demethylation could reactivate *P16* gene transcription, which subsequently inhibited the migration and growth of cancer cells *in vivo*.

It is well known that an appropriate proportion of 5hmCs in the genome is distributed with a strand bias [9,28]. We recently reported that there were dense 5hmCs in the *P16* exon-1 AS-strands in HCT116 cells [21,22]. Based on the comprehensive TAB-DHPLC and TAB sequencing results, here, we further demonstrated that 5hmCs were enriched only in the AS-strand of the *P16* exon-1 in HCT116 cells and AGS cells, while sporadic 5hmCs were detected in the S-strand of *P16* promoter and exon-1 regions. The fact that 88% of the exon-1 AS-strand CpGs in the wild-type *P16* alleles in HCT116 cells are hydroxymethylated indicates that *P16* exon-1 has a methylation: hydroxymethylation (M:H) mixture, composed of a fully hydroxymethylated AS-strand and a truly methylated S-strand.

It has been reported that triple knockout of *TET-1/2/3* led to bivalent promoter hypermethylation in H1 cells [33]. TSSs are DNA replication start sites. S- and AS-strands of genes are generally replicated by different types of DNA polymerases in eukaryotic cells (Polδ for the leading strand and Polα for the lagging strand). Unlike true DNA methylation that is maintained by DNMT1 during DNA synthesis in the S-phase of the cell cycle, DNA hydroxymethylations are probably maintained by the *de novo* methyltransferases DNMT3a/b [34]. It is of great interest to study the mechanisms leading to the strand bias of DNA hydroxymethylation.

Three types of epigenetic editing methods, including ZFP-, transcription activator-like effector (TALE)-, and CRISPR/dCas9-based systems, have emerged as advanced tools to study the functions of epigenetic modifications [19,35–39]. According to the reported data, the specificity and efficiency of ZFP-based epigenetic editing tools are likely higher than those of TALE-based editing tools or CRISPR/dCas9-based editing tools. For example, the expression controllable ZFP-based P16-Dnmt could selectively methylate entire *P16* CGIs around the TSS [19]. However, CRISPR/dCas9-Dnmt3a, combined with *P16-*sgRNA, could specifically methylate only approximately 50 bp sgRNA target-flanking sequences (not including the sgRNA target) [40,41]. In contrast, *P16* TALE-Dnmt could methylate the *P16* target and other CGIs within the *P15-P14-P16-MTAP* gene cluster and repress their transcription, with low specificity [42]. We recently reported that *ANRIL* expression was repressed in cancer cells by *P16* methylation [20]. Here, we further demonstrated that the *ANRIL* expression was upregulated in the P16-TET-expressing cells and that the mRNA levels of *P15* and *P14* were not increased. These observations suggest that P16-TET could specifically demethylate *P16* CGIs via DNA hydroxymethylation and reactivate the transcription of both the *P16* and *ANRIL* genes, though DNA hydroxymethylation itself could not reactivate *P16* gene transcription.

Recently, we found that all *P16* mRNA clones in the HCT116 cells were transcribed only from the unmethylated *P16* alleles, and none from the methylated: hydroxymethylated (M:H) *P16* alleles [21], and that both true P16M and P16H could similarly increase the risk for malignant transformation of oral epithelial dysplasia in a prospective study [23]. The findings of the present study show that the P16-TET-induced hydroxymethylation of *P16* alleles in both H1299 and AGS cells retain transcriptional silence, which provides a possible mechanism to explain the above observations.

There are many differences between cell culture and animal models. Although the proliferation of H1299 cells that are stably transfected with P16-TET was not changed under *in vitro* culture conditions, the growth of xenograft tumors from these cells was obviously inhibited in host mice. The exact reasons leading to this difference are unknown; however, the reactivation of methylated *P16* alleles via DNA demethylation by P16-TET may account for the growth inhibition *in vivo*. The growth inhibition of xenograft tumors from the *P16-*deleted A549 control cells was not observed, suggesting that the growth inhibition of xenografts by P16-TET may be a *P16-*dependent phenomenon. In the rescue assay, siRNA knockdown of P16-TET-reactivated *P16* expression almost completely reversed the inhibition of P16-TET-induced cell migration. This further suggests that the inhibition of the cancer cell migration by P16-TET may be a *P16-*specific effect.

In conclusion, we found that hydroxymethylation of *P16* CGI is located mainly in the exon-1 AS-strand. P16H alleles are transcriptionally inactive. *P16* demethylation via hydroxymethylation could reactivate gene transcription and inhibit the growth of cancer cells.

## METHODS

### Cell Lines and Culture

The colon cancer cell line HCT116 was purchased from the American Type Culture Collection (ATCC). The GC cell line AGS and the lung cancer cell line H1299 were kindly provided by Prof. Chengchao Shou from the Peking University Cancer Hospital and Institute. The colon cancer cell line, RKO was kindly provided by Prof. Guoren Deng from the University of California, San Francisco. These cells were cultured in RPMI 1640 containing 10% FBS and 100 U/mL penicillin/streptomycin (Invitrogen, California, USA) at 37°C in a humidified incubator with 5% CO_2_.

These cell lines were tested and authenticated by Beijing JianLian Genes Technology Co., LTD before they were used in this study. STR patterns were analyzed using a Goldeneye™20A STR Identifiler PCR Amplification Kit. Gene Mapper v3.2 software (ABI) was used to match the STR pattern with the ATCC online databases.

### Characterization of 5mC and 5hmC Sites in *P16* CGIs

Total P16M was analyzed using 150-bp regular methylation-specific PCR (MSP) [43]. To selectively detect P16H, the genomic DNA (3 μg), spiked with *M.sss*I-methylated and 5hmC-containing λ-DNA controls, was modified using the TET-Assisted Bisulfite (TAB) Kit, according to manufacturer’s instructions (WiseGene, Cat# K001). During TAB-modification, 5mC was oxidized to 5caC, and both 5caC and unmethylated cytosine were subsequently converted to uracil through bisulfite-induced deamination, whereas 5hmC was protected from oxidation via 5hmC-specific β-glucosylation [25]. The conversion rates of unmethylated cytosine, 5mCs, and 5hmCs in the bisulfite-/TAB-treated λ-DNA controls were 100%, 99.7%, and 1.5%, respectively (Figure S1). P16H was analyzed using the TAB-MSP.

The proportion of hydroxymethylated S- and AS-strands of the *P16* promoter and exon-1 CGIs were analyzed using DHPLC and clone sequencing, respectively [45,46]. The adjusted ratio of the peak height for the hydroxymethylated region to that of the unmethylated region was used to represent the P16H proportion that was adjusted. The ratio of the P16M peak height to the P16U peak height for *P16-*hemimethylated HCT116 cells was used as a reference. The sequences of the universal primers used to amplify these fragments are listed in Table 1.

**Table 1.**
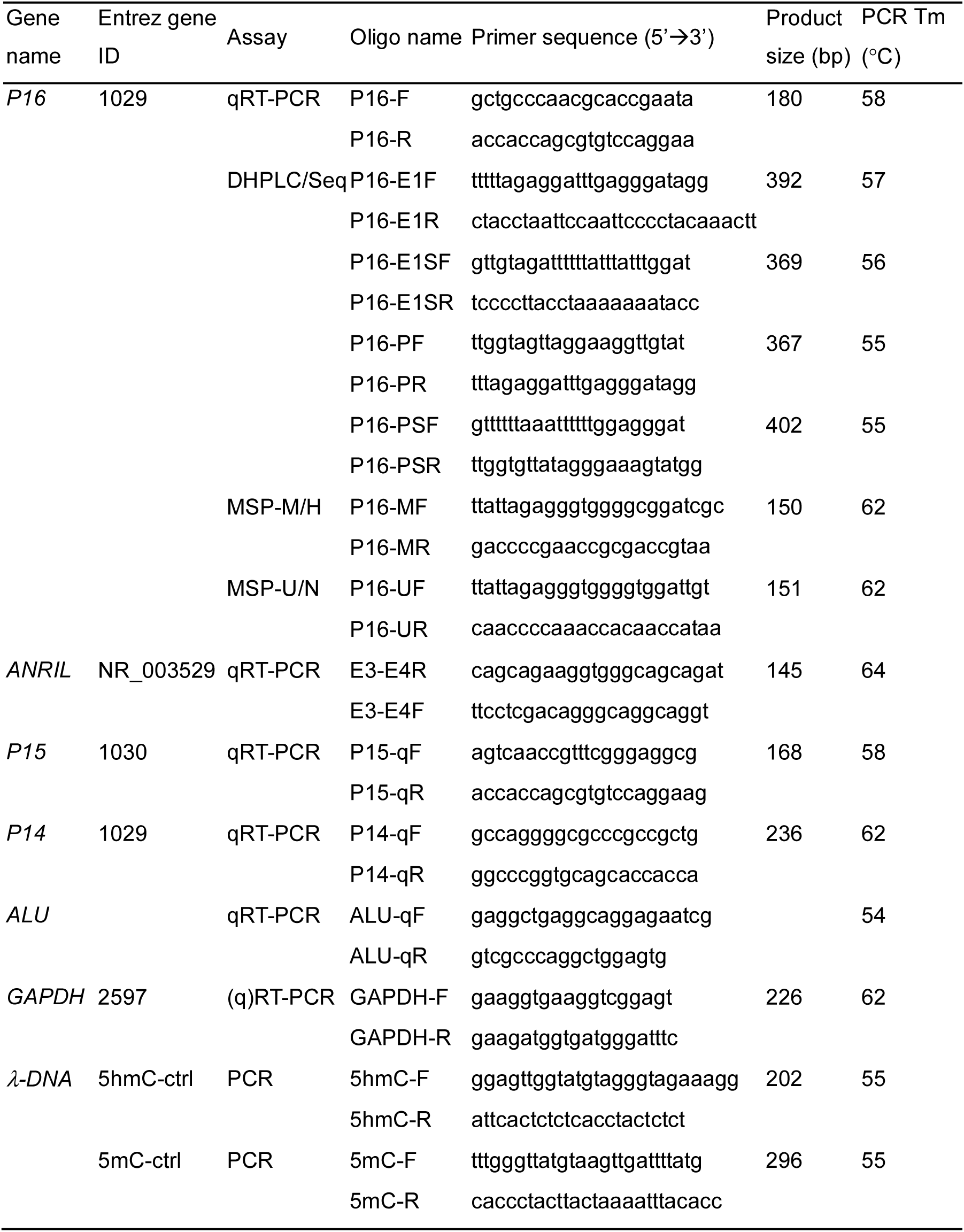
Sequences of oligonucleotides used as primers in various PCR-based assays

### Construction of Expression Vectors and Transfection

To construct the *P16-*specific DNA dioxygenase (P16-TET) expression vector, an SP1-like engineered seven-zinc finger protein (7ZFP-6I) [19] that can specifically bind to the 21-bp fragment (5’-gaggaaggaaac<u>ggggcgggg</u>-3’, including an <u>Sp1-binding site</u>) within the human *P16* core promoter [24,47], was fused with the catalytic domain (CD: 1418-2136 aa) of human *TET1* (NM_030625.2) [48] and inserted into a pcDNA3.1b vector and then used in transient transfection assays. An inactive P16-TET mutant containing an H1671Y mutation in the CD domain vector was also constructed and used as a negative control vector (Figure S2A). The P16-TET sequence was further integrated into the expression-controllable pTRIPZ vector carrying a “Tet-on” switch (Open Biosystem, USA) (Figure S2B) [19]. Purified P16-TET pTRIPZ plasmid was mixed with VSVG and Δ8.9 (Addgene, USA) to prepare lentivirus transfection particles. The fresh lentivirus particles were used to stably infect AGS and H1299 cells containing homogenously methylated *P16* CpG islands. Doxycycline (Dox; final conc. 0.25 μg/mL) was added to the medium to induce P16-TET expression.

Two *P16-*specific siRNAs (5’-ccgua aaugu ccauu uauatt-3’ and 5’–uauaa augga cauuu acggtt-3’) were synthesized (GenePharma, Shanghai) and used to transiently transfect cells at a final concentration of 1.0 μg/mL. Two scrambled siRNAs (5’-uucuc cgaac guguc acgutt-3’ and 5’-acgug acacg uucgg agaatt-3’) were used as negative controls (NC).

### Treatment of 5’-Aza-Deoxycytidine (DAC)

The AGS cells were treated with DAC (final concentration 20 nM; Abcam ab120842, Cambridge, UK) for 7 days in the P16-immunostaining assay or 10 days prior to FACS sorting.

### Extraction of RNA and Quantitative RT-PCR (qRT-PCR)

Cells were harvested when they reached a confluency of approximately 70%. Total RNA was extracted by TRIzol (Invitrogen, California, USA). The cDNA was reverse-transcribed using the ImProm-II™ Reverse Transcription System (A3800; Promega). The expression levels of the *ANRIL*, *P16*, *P15*, *P14*, and *TET-1/2/3* genes were analyzed by quantitative RT-PCR using the corresponding primer sets (Table 1), as previously described [20]. Power SYBR Green PCR Master Mix (Fermentas, Canada) was used in the qRT-PCR analyses (ABI-7500FAST). The relative mRNA level was calculated based on the average Ct value of the target gene and the *Alu* reference [2^-(Ct_target_gene_-Ct_Alu_)^] [49].

### Western Blot and Confocal Microscopy Analysis of the P16 Expression Status

The *P16* mRNA and protein levels in the cells were analyzed as previously described [19]. Rabbit monoclonal antibody against human P16 protein (ab108349, Abcam, Britain) was used in the Western blot assay, and mouse monoclonal antibody against the human P16 protein (Ventana Roche-E6H4, USA) was used in the immunostaining assay.

### Cell FACS Sorting

The P16-TET stably transfected H1299 cells (treated with doxycycline for 21 days) and AGS cells (treated with 5-aza-deoxycytidine for 10 days) were fixed with methanol, permeabilized with 0.1% Tween-20 in PBS, pretreated with 10% fetal bovine serum and 0.3 M glycine in PBS, and were then stained with the mouse monoclonal antibody against the human P16 protein (Ventana Roche-E6H4, USA) and the FITC-tagged secondary antibody. The P16-staining cell population proportion was determined using an immuno-fluorescence confocal microscope. These cells were sorted by FACS and divided into three subpopulations, strong-, weak-, and non-P16-staining, using P16-TET H1299 cells without doxycycline treatment or AGS cells without DAC treatment as P16 protein negative controls. According to the confocal analysis results, we setup the cutoff value to sort definite and indefinite P16 protein positive (P16(+) and P16(±)) cell subpopulations. The strong and weak FITC-staining cells were called as the P16(+) and P16(±) subpopulations, respectively.

### IncuCyte ZOOM and Transwell Migration Tests

The long-term live content kinetic imaging platform (IncuCyte Zoom, Essen BioSci, USA) was used to dynamically detect the proliferation and migration of live cancer cells. The phase object confluence (%) was used to generate a cell proliferation curve. The relative wound density, a measure (%) of the density of the wound region relative to the density of the cell region, was used as the metric for cell migration. The transwell migration test was performed as previously described [19].

### Xenografts in SCID Mice

Cells stably transfected with the P16-TET vector were induced with 0.25 μg/mL doxycycline for 7 days and then subcutaneously injected into one lower limb of each NOD-SCID mouse (10^5^ cells/injection; female, 5 weeks old, 10∼20 g, purchased from Beijing Huafukang Biotech). The negative control cells stably transfected with the empty pTRIPZ vector were simultaneously injected into the opposite side of each mouse. These mice were given distilled, sterile water containing 2 μg/mL doxycycline and were sacrificed on the 50^th^ post-transplantation day. The xenografts were weighed and histologically confirmed [19]. Two repeat experiments were performed.

### Statistical Analysis

Student’s t-test was used for statistical analysis. All *P*-values were two-sided, and a *P-*value of <0.05 was considered to be statistically significant.

## Author contributions

YG and PL: elucidated the biological function of *P16* hydroxymethylation; SQ: discovered *P16* hydroxymethylation and its association with gastric carcinogenesis; XH: demonstrated the strand-bias distribution of 5hmCs in the *P16* alleles; CC, Z-mL, and BZ: constructed the *P16-*specific oxygenase; LG performed the animal experiments; BZ performed immunostaining and cell sorting assays and codesigned the study; ZL and JZ: carried out other experiments; DD: designed the study, analyzed the data, and wrote the manuscript. All authors read and approved the final manuscript.

## ACKNOWLEDGEMENTS

We thank Mr. Jordan M. Grainger (Predoctoral student, Molecular Pharmacology and Experimental Therapeutics, Mayo Clinic, Rochester, Minnesota, USA) and Dr. Huidong Shi (Cancer Center, Georgia Medical College, Augusta, USA) for English language editing.

## Ethics approval and consent to participate

The Peking University Cancer Hospital and Institutional Review Boards approved this study. Ethical approval for the animal experiments was obtained.

## Disclosure of potential conflicts of interest

The authors have declared that no competing interests exist.

## SUPPORTING FIGURE LEGENDS

**Figure S1.**
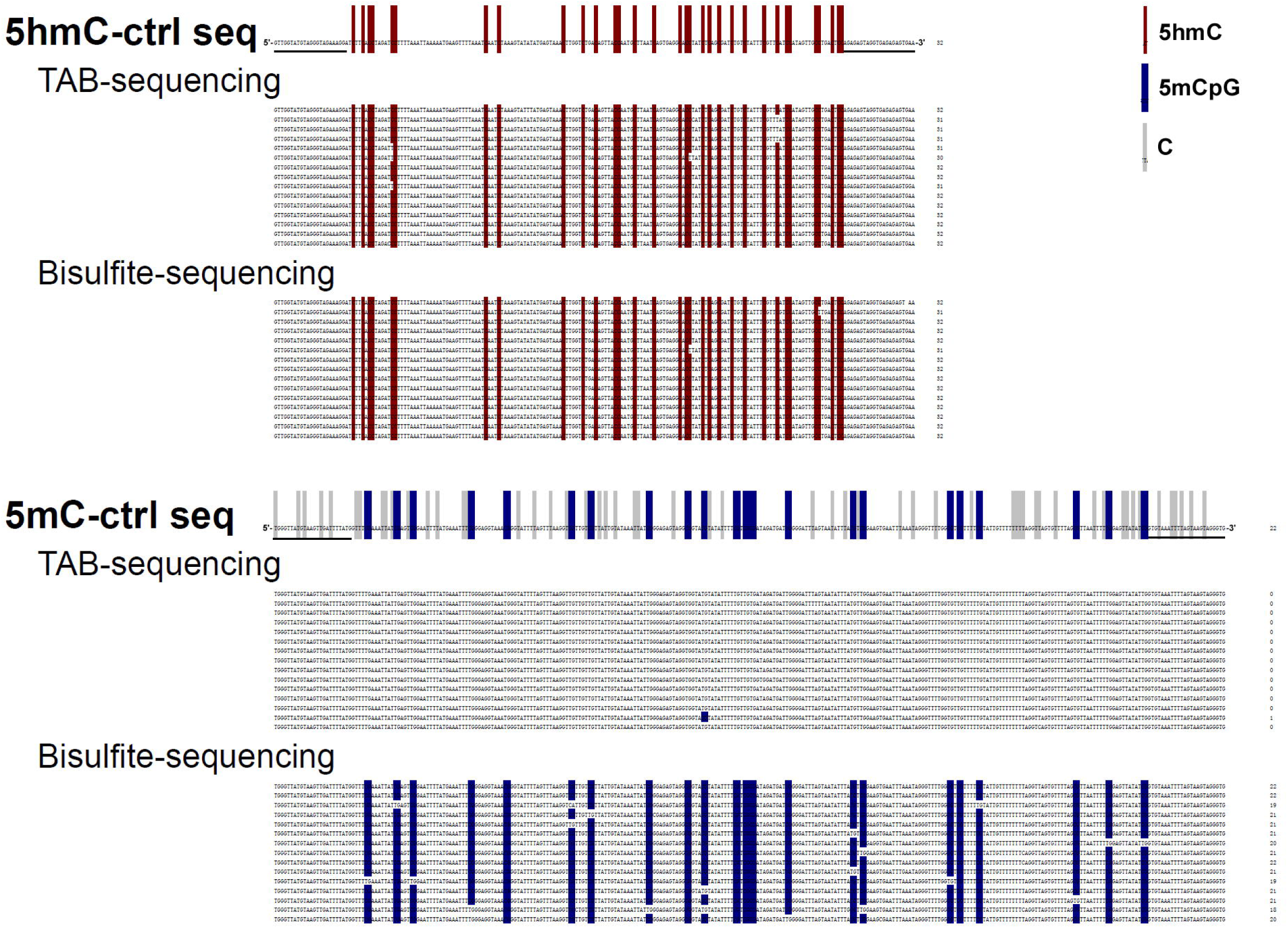
Characterization of the true methylation and hydroxymethylation states of CpG sites in the *M.sss*I-methylated and 5hmC-containing λ-DNA controls (5mC-ctrl and 5hmC-ctrl). Bisulfite-modified DNA templates were used to discriminate 5mC or 5hmC from unmethylated cytosine. TAB-modified DNA templates were used to discriminate 5hmC from 5mC and unmethylated cytosine. The CpG sites within the consensus sequences are listed above the corresponding clone sequences. The number of 5hmC and 5mC sites within each clone is also listed on the right side. The control DNA was added into the test samples to monitor the conversion status of 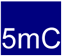, 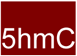, and 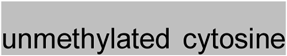 in genomic DNA by bisulfite and TAB treatments.

**Figure S2.**
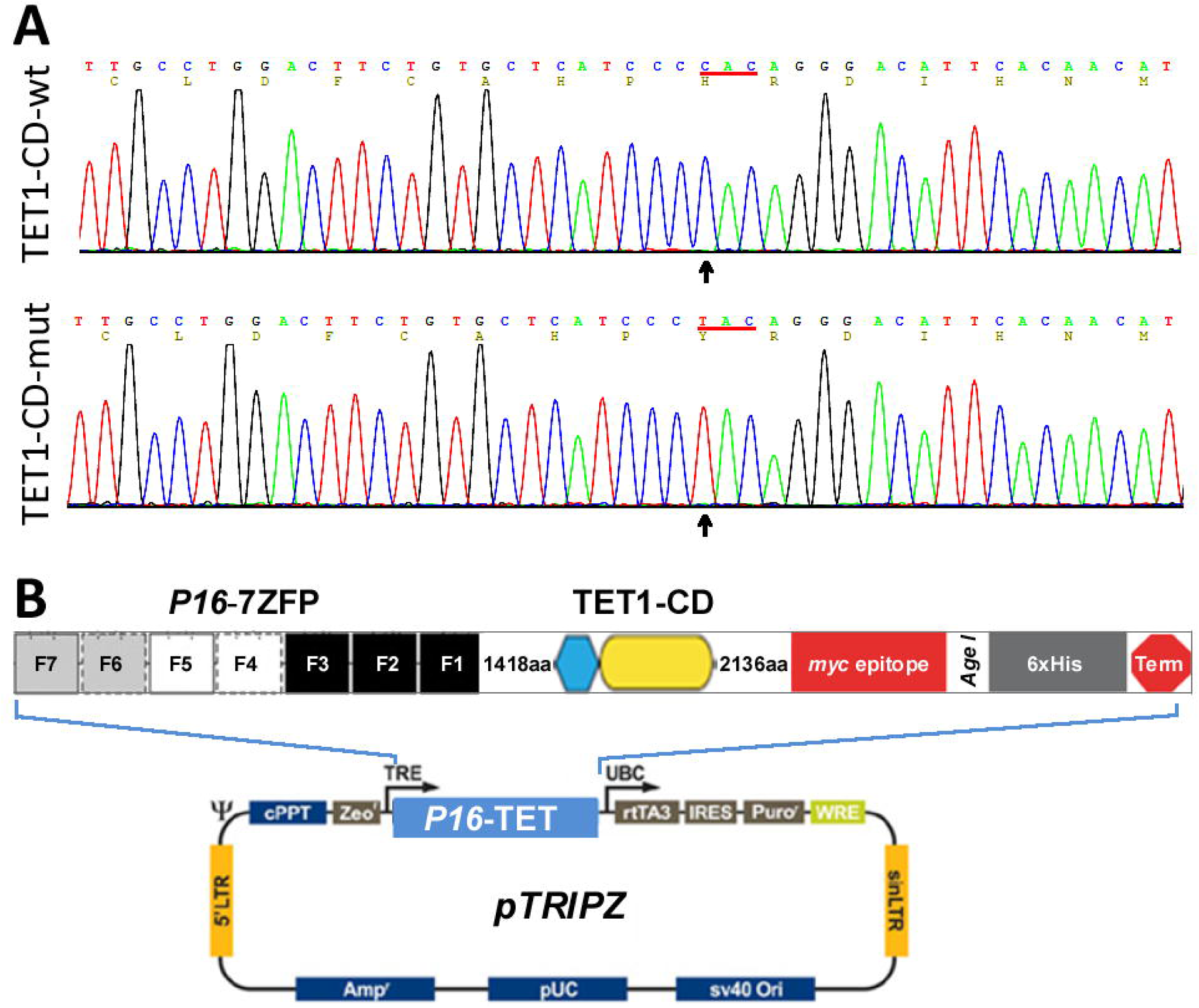
Construction of P16-TET expression vector. (**A**) Fragment sequences of the catalytic domain (CD) of the human *TET1* gene were used to construct the wild-type P16-TET and its inactive H1671Y-mutant control. (**B**) The pTRIPZ vector integrated with 7ZFP-6I and the TET1 CD domain.

**Figure S3.**
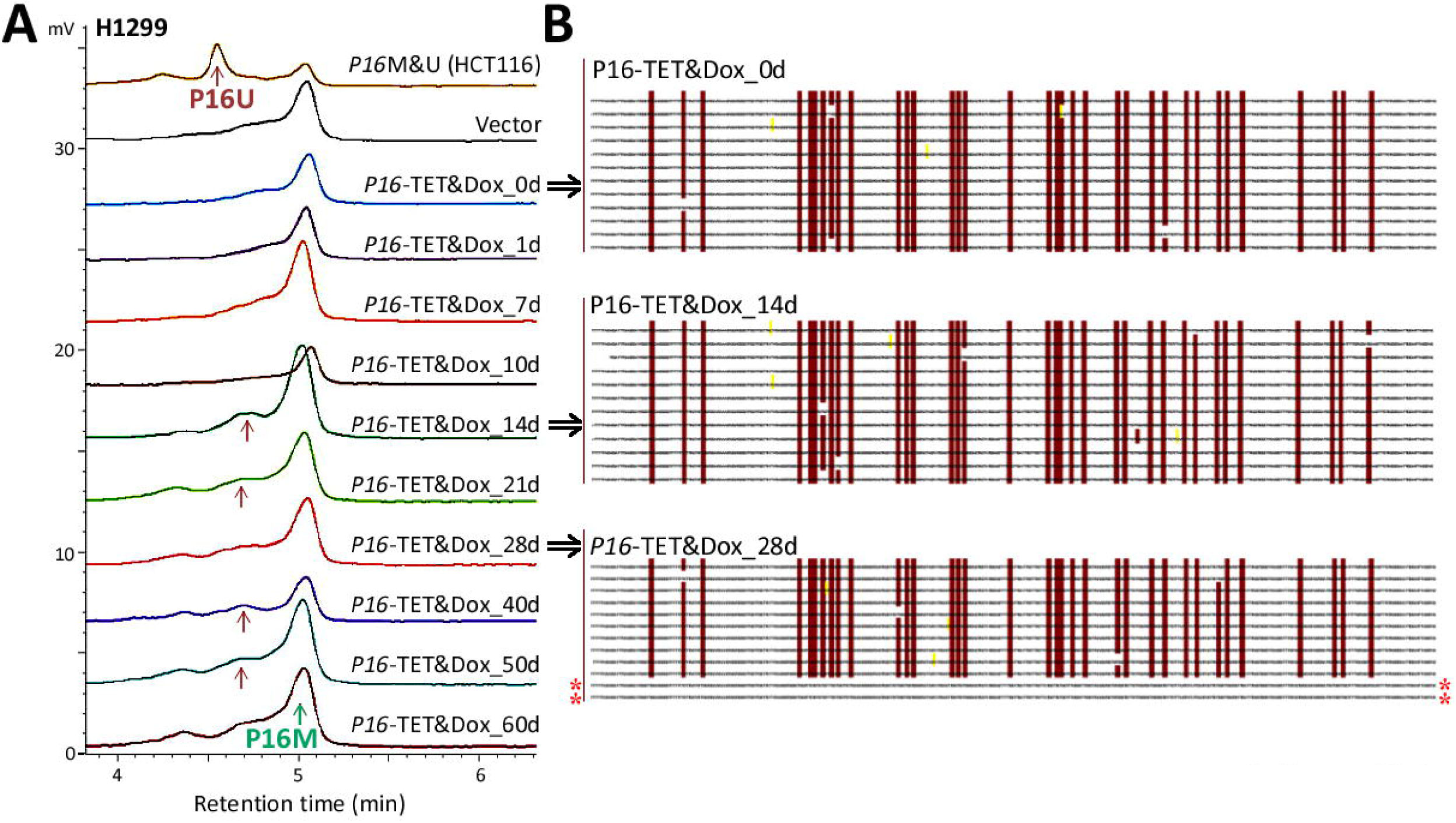
The demethylation status of *P16* CpG islands and reactivation of *P16* expression in H1299 cells. (**A**) Bisulfite-DHPLC analysis for detecting methylated- and demethylated-*P16* (P16-M and P16-U) in H1299 cells stably transfected with P16-TET or empty control vector after doxycycline treatment for different days; genomic DNA from HCT116 cells was used as a P16-M and P16-U control. (**B**) Bisulfite sequencing analysis for detecting the methylation status of *P16* exon-1 antisense strands from H1299 cells stably transfected with P16-TET and doxycycline-treated for 0, 14, and 28 days.

**Figure S4.**
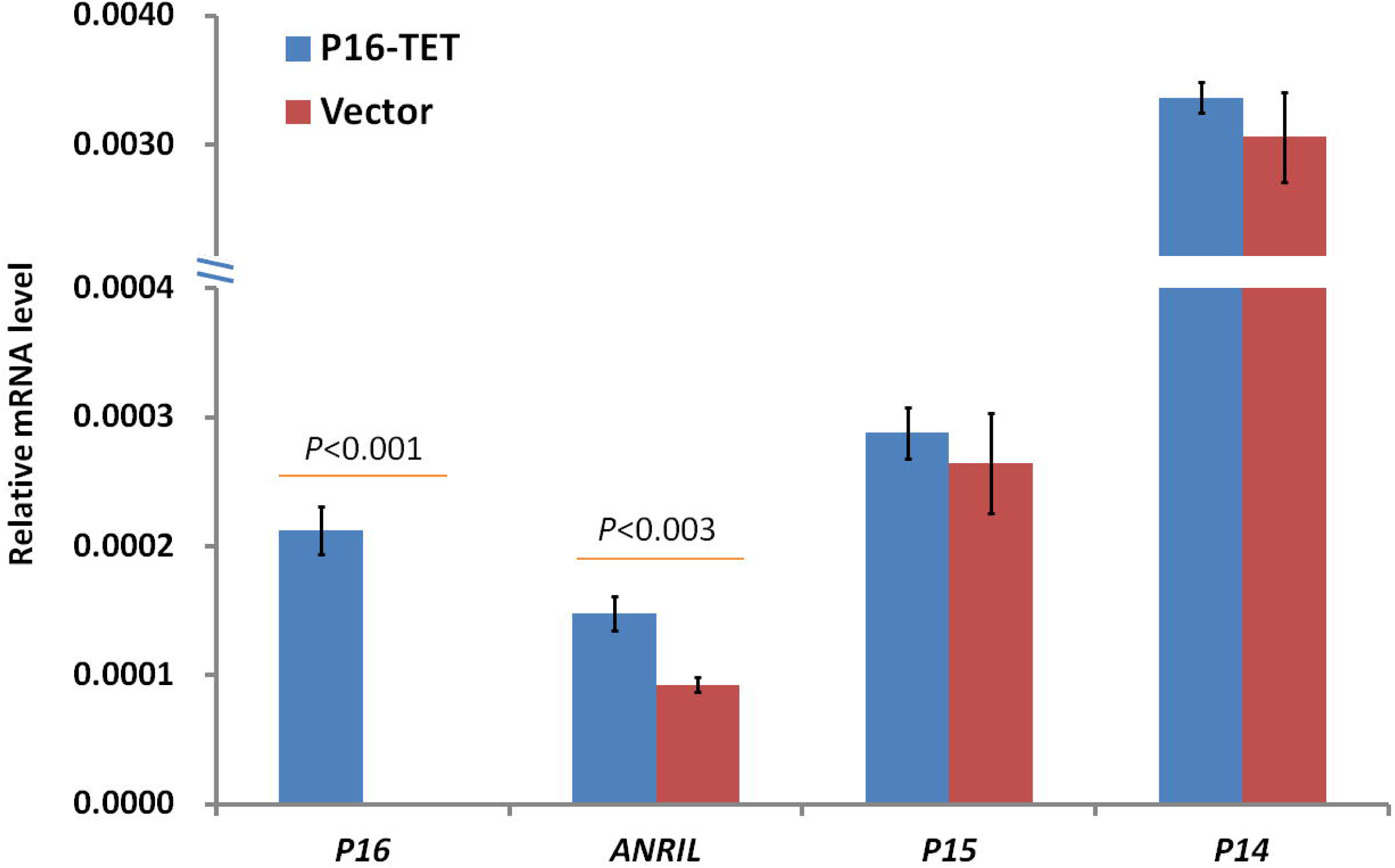
Effect of P16-TET on transcription of *P16*, *ANRIL*, *P15*, and *P14* genes in the 9p21 locus of H1299 cells stably transfected with the P16-TET pTRIPZ vector and treated with doxycycline for 14 days.

**Figure S5.**
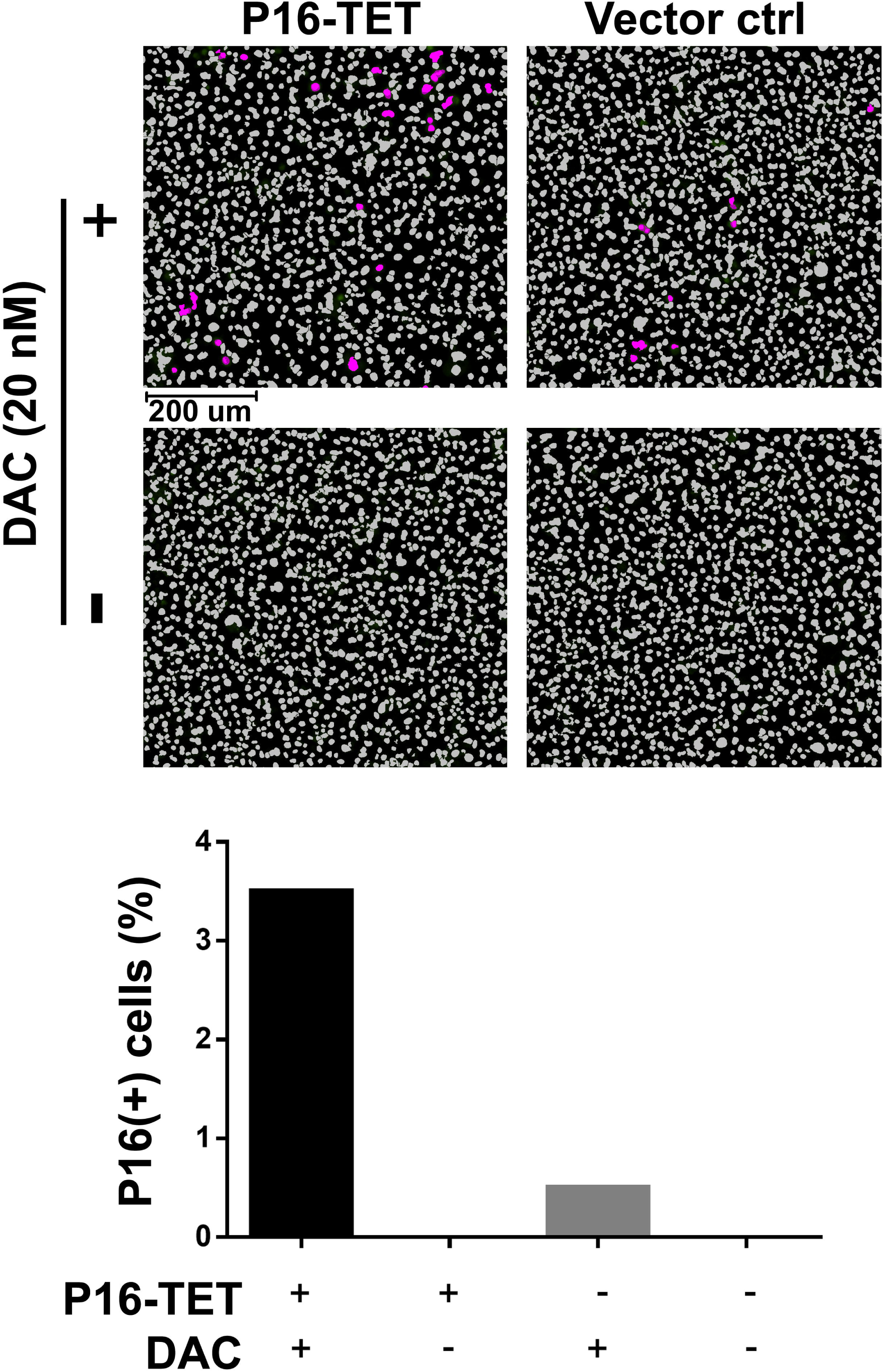
P16 expression status of *P16-*methylated AGS cells stably transfected with P16-TET (or control vector) as shown by in immunostaining. P16-TET cells were cotreated with 5-aza-deoxycytidine (DAC, final concentration 20 nM) or its reagent control for 10 days (without doxycycline treatment). The proportion of P16(+) cells in the confocal microscopy images for each group was automatically counted using the ImageXpress Micro High Content Screening System (Molecular Devices, USA).

**Figure S6.**
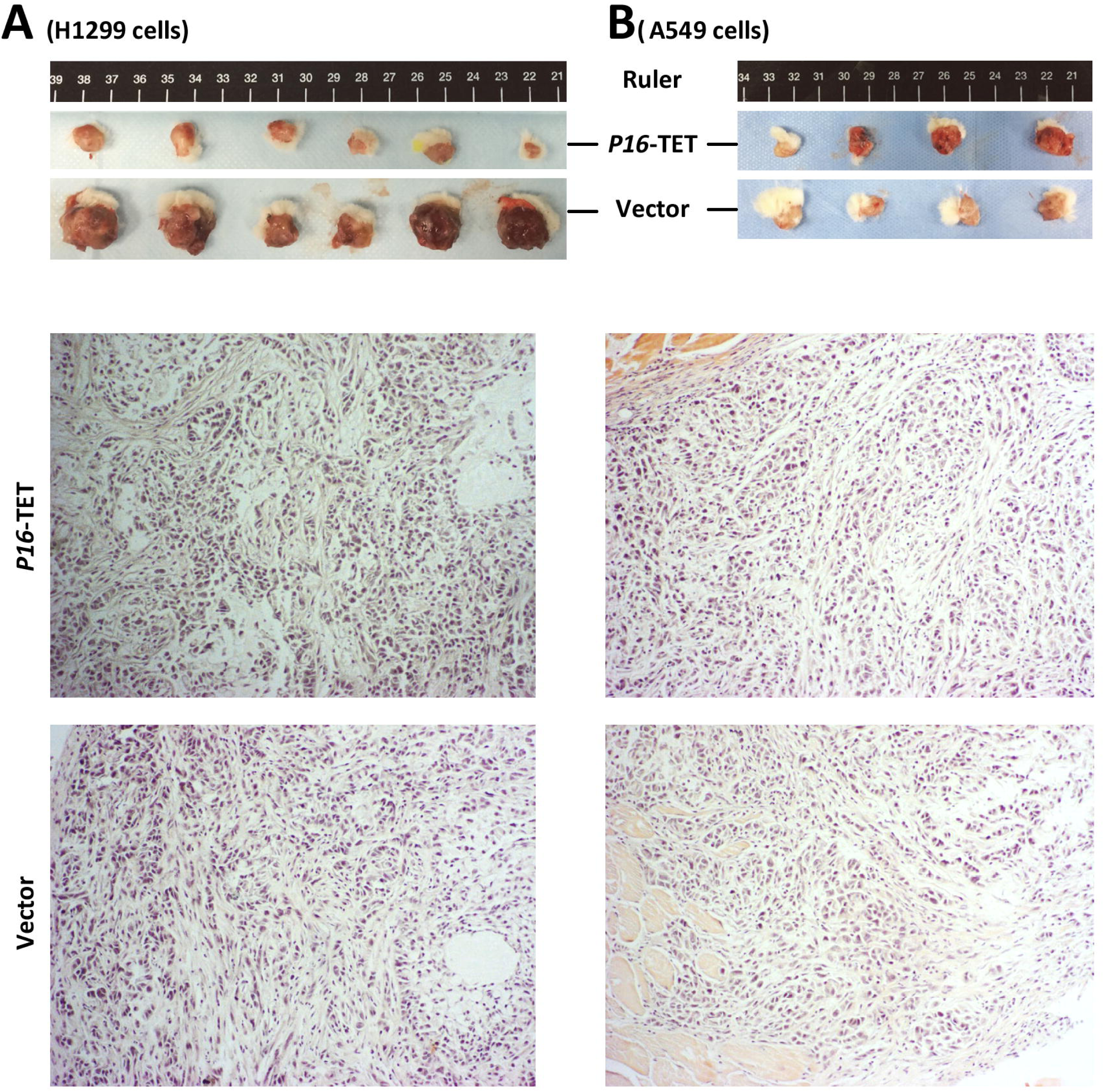
Effects of P16-H on the growth of xenograft tumors from H1299 and A549 cells in NOD-SCID mice. (**A**) Images of xenograft tumors from H1299 cells with and without stable P16-TET transfection on the 36^th^ experimental day. (**B**) Images of xenograft tumors from *ink4a/b*-deleted A549 cells with and without stable P16-TET transfection on the 33^rd^ experimental day. The H.E. staining images are also displayed.

**Figure S7.**
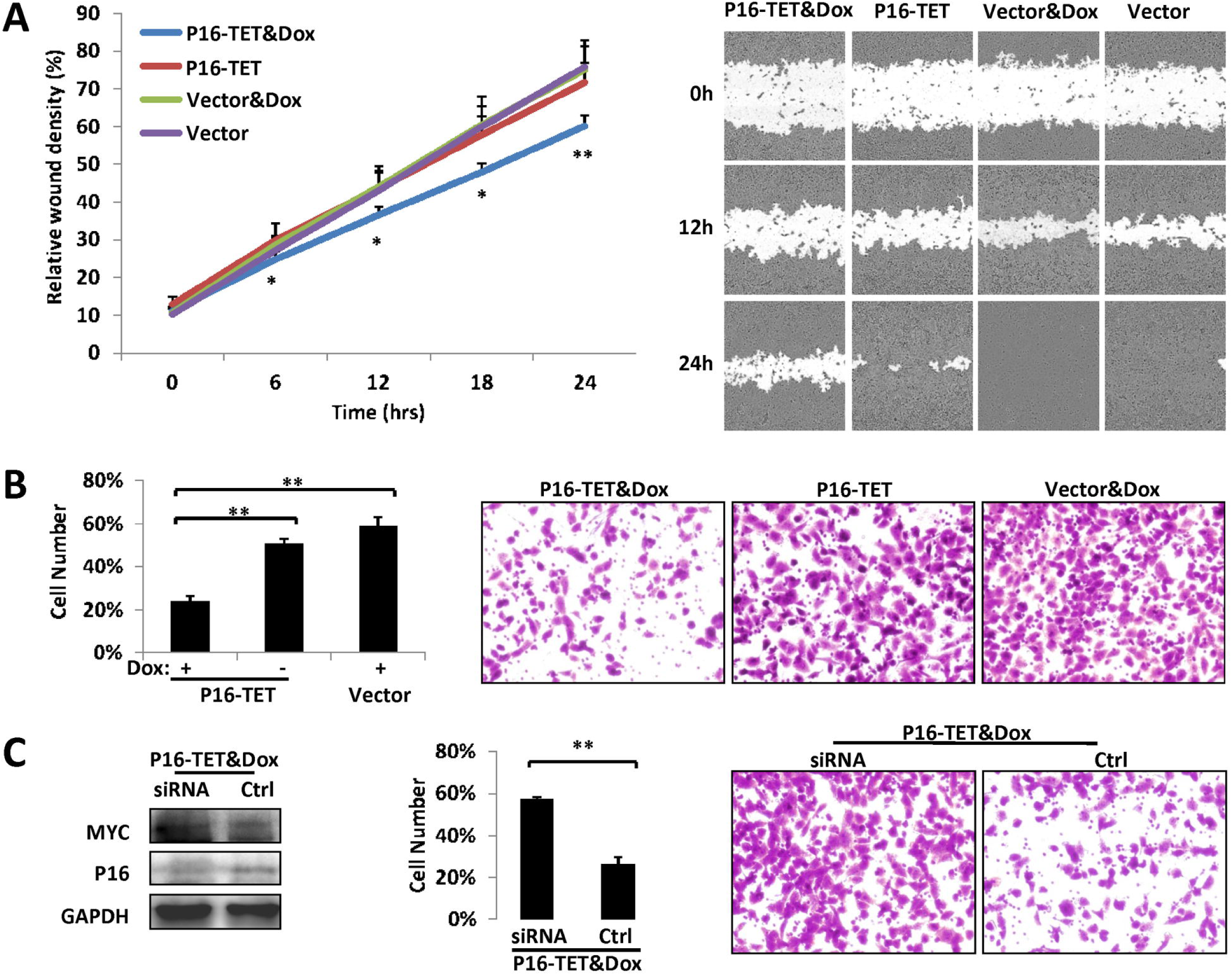
Effects of *P16* expression changes induced by P16-TET and *P16* siRNA on the migration of H1299 cells stably transfected with P16-TET *in vitro*. (**A**) IncuCyte ZOOM scratch assay for detecting cell migration. (**B**) Transwell migration assay for detecting the migration of cells with 24 hr incubation. The average cell number (confluence) and s.d. value are displayed in the left charts. (**C**) Rescue assay for detecting the effect of siRNA-knockdown of *P16* expression on the migration of H1299 cells stably transfected with P16-TET and treated with doxycycline for 14 days. The cells (4.5×10^4^) transiently transfected with two types of *P16*-specific siRNAs were seeded into each well and incubated for 72 hrs. The expression status of the P16 protein was monitored using Western blot assay.

## REFERENCES

[1] Tahiliani M, Koh KP, Shen Y, Pastor WA, Bandukwala H, Brudno Y, et al. Conversion of 5-methylcytosine to 5-hydroxymethylcytosine in mammalian DNA by MLL partner TET1. Science 2009; 324: 930–935.

[2] Kriaucionis S, Heintz N. The nuclear DNA base 5-hydroxymethylcytosine is present in Purkinje neurons and the brain. Science 2009; 324: 929–930.

[3] He YF, Li BZ, Li Z, Liu P, Wang Y, Tang Q, et al. Tet-mediated formation of 5-carboxylcytosine and its excision by TDG in mammalian DNA. Science 2011; 333: 1303–1307.

[4] Ito S, Shen L, Dai Q, Wu SC, Collins LB, Swenberg JA, et al. Tet proteins can convert 5-methylcytosine to 5-formylcytosine and 5-carboxylcytosine. Science 2011; 333: 1300–1303.

[5] Gu TP, Guo F, Yang H, Wu HP, Xu GF, Liu W, et al. The role of Tet3 DNA dioxygenase in epigenetic reprogramming by oocytes. Nature 2011; 477: 606–610.

[6] Hackett JA, Sengupta R, Zylicz JJ, Murakami K, Lee C, Down TA, et al. Germline DNA demethylation dynamics and imprint erasure through 5-hydroxymethylcytosine. Science 2013; 339: 448–452.

[7] Pastor WA, Pape UJ, Huang Y, Henderson HR, Lister R, Ko M, et al. Genome-wide mapping of 5-hydroxymethylcytosine in embryonic stem cells. Nature 2011;473:394–7

[8] Ficz G, Branco MR, Seisenberger S, Santos F, Krueger F, Hore TA, et al. Dynamic regulation of 5-hydroxymethylcytosine in mouse ES cells and during differentiation. Nature 2011; 473: 398–402.

[9] Yu M, Hon GC, Szulwach KE, Song CX, Zhang L, Kim A, et al. Base-resolution analysis of 5-hydroxymethylcytosine in the mammalian genome. Cell 2012; 149: 1368–1380.

[10] Merlo A, Herman JG, Mao L, Lee DJ, Gabrielson E, Burger PC, et al. 5’ CpG island methylation is associated with transcriptional silencing of the tumor-suppressor p16/cdkn2/mts1 in human cancers. Nature Medicine 1995; 1: 686–692.

[11] Herman JG, Merlo A, Mao L, Lapidus RG, Issa JPJ, Davidson NE, et al. Inactivation of the cdkn2/p16/mts1 gene is frequently associated with aberrant dna methylation in all common human cancers. Cancer Research 1995; 55: 4525–4530.

[12] Sun Y, Deng DJ, You WC, Bai H, Zhang L, Zhou J, et al. Methylation of p16 CpG islands associated with malignant transformation of gastric dysplasia in a population-based study. Clinical Cancer Research 2004; 10: 5087–5093.

[13] Belinsky SA, Liechty KC, Gentry FD, Wolf HJ, Rogers J, Vu K, et al. Promoter hypermethylation of multiple genes in sputum precedes lung cancer incidence in a high-risk cohort. Cancer Res 2006; 66: 3338–3344.

[14] Hall GL, Shaw RJ, Field EA, Rogers SN, Sutton DN, Woolgar JA, et al. p16 Promoter methylation is a potential predictor of malignant transformation in oral epithelial dysplasia. Cancer Epidemiol Biomarkers Prev 2008; 17: 2174–2179.

[15] Cao J, Zhou J, Gao Y, Gu L, Meng H, Liu H, et al. Methylation of p16 CpG Island Associated with Malignant Progression of Oral Epithelial Dysplasia: A Prospective Cohort Study. Clinical Cancer Research 2009; 15: 5178–5183.

[16] Jin Z, Cheng Y, Gu W, Zheng Y, Sato F, Mori Y, et al. A multicenter, double-blinded validation study of methylation biomarkers for progression prediction in Barrett’s esophagus. Cancer Res 2009; 69: 4112–4115.

[17] Liu HW, Liu XW, Dong GY, Zhou J, Liu Y, Gao Y, et al. *P16* Methylation as an Early Predictor for Cancer Development From Oral Epithelial Dysplasia: A Double-blind Multicentre Prospective Study. EBioMedicine 2015; 2: 432–437.

[18] Gao H-e, Zhang Y, Zhou J, Li Z, Ma J-l, Liu W-d, et al. Association between p16 methylation and malignant transformation of gastric dysplasia. Chinese Journal of Cancer Prevention and Treatment 2017; 24: 431–436.

[19] Cui C, Gan Y, Gu L, Wilson J, Liu Z, Zhang B, et al. P16-specific DNA methylation by engineered zinc finger methyltransferase inactivates gene transcription and promotes cancer metastasis. Genome Biology 2015; 16: 252.

[20] Gan Y, Ma W, Wang X, Qiao J, Zhang B, Cui C, et al. Coordinate Transcription of A*NRIL* and P*16*Genes Silenced by DNA Methylation. Chinese Journal of Cancer Research 2018; 30: 93–103.

[21] Qin SS, Li Q, Zhou J, Liu ZJ, Su N, Wilson J, et al. Homeostatic Maintenance of Allele-Specific p16 Methylation in Cancer Cells Accompanied by Dynamic Focal Methylation and Hydroxymethylation. Plos One 2014; 9: E97785.

[22] Qin S, Zhang B, Tian W, Gu L, Lu Z, Deng D. Kaiso mainly locates in the nucleus in vivo and binds to methylated, but not hydroxymethylated DNA. Chinese Journal of Cancer Research 2015; 27: 148–155.

[23] Liu HW, Liu ZJ, Liu XW, Xu S, Wang L, Liu Y, et al. A similar effect of *P16* hydroxymethylation and true-methylation on the prediction of malignant transformation of oral epithelial dysplasia: observation from a prospective study. BMC Cancer 2018; 18:918.

[24] Zhang B, Xiang S, Zhong Q, Yin Y, Gu L, Deng D. The p16-Specific Reactivation and Inhibition of Cell Migration Through Demethylation of CpG Islands by Engineered Transcription Factors. Human Gene Therapy 2012; 23: 1071–1081.

[25] Sun Z, Terragni J, Jolyon T, Borgaro JG, Liu Y, Yu L, et al. High-resolution enzymatic mapping of genomic 5-hydroxymethylcytosine in mouse embryonic stem cells. Cell Rep 2013; 3: 567–576.

[26] Wang T, Wu H, Li Y, Szulwach KE, Lin L, Li X, et al. Subtelomeric hotspots of aberrant 5-hydroxymethylcytosine-mediated epigenetic modifications during reprogramming to pluripotency. Nat Cell Biol 2013; 15: 700–711.

[27] Kim M, Park YK, Kang TW, Lee SH, Rhee YH, Park JL, et al. Dynamic changes in DNA methylation and hydroxymethylation when hES cells undergo differentiation toward a neuronal lineage. Hum Mol Genet 2014; 23: 657–667.

[28] Wen L, Li X, Yan L, Tan Y, Li R, Zhao Y, et al. Whole-genome analysis of 5-hydroxymethylcytosine and 5-methylcytosine at base resolution in the human brain. Genome Biol 2014; 15: R49.

[29] Chen K, Zhang J, Guo Z, Ma Q, Xu Z, Zhou Y, et al. Loss of 5-hydroxymethylcytosine is linked to gene body hypermethylation in kidney cancer. Cell Res 2016; 26: 103–118.

[30] Li X, Liu Y, Salz T, Hansen KD, Feinberg A. Whole-genome analysis of the methylome and hydroxymethylome in normal and malignant lung and liver. Genome Res 2016; 26: 1730–1741.

[31] Huang Y, Chavez L, Chang X, Wang X, Pastor WA, Kang J, et al. Distinct roles of the methylcytosine oxidases Tet1 and Tet2 in mouse embryonic stem cells. Proc Natl Acad Sci U S A 2014; 111: 1361–1366.

[32] Uribe-Lewis S, Stark R, Carroll T, Dunning MJ, Bachman M, Ito Y, et al. 5-hydroxymethylcytosine marks promoters in colon that resist DNA hypermethylation in cancer. Genome Biol 2015; 16: 69.

[33] Verma N, Pan H, Doré LC, Shukla A, Li QV, Pelham-Webb B, et al. TET proteins safeguard bivalent promoters from de novo methylation in human embryonic stem cells. Nat Genet 2018; 50: 83–95.

[34] Hashimoto H, Liu Y, Upadhyay AK, Chang Y, Howerton SB, Vertino PM, et al. Recognition and potential mechanisms for replication and erasure of cytosine hydroxymethylation. Nucleic Acids Res 2012; 40: 4841–4849.

[35] Liu XS, Wu H, Ji X, Stelzer Y, Wu X, Czauderna S, et al. Editing DNA Methylation in the Mammalian Genome. Cell 2016; 167: 233–247.

[36] Kearns NA, Pham H, Tabak B, Genga RM, Silverstein NJ, Garber M, et al. Functional annotation of native enhancers with a Cas9-histone demethylase fusion. Nat Methods 2015; 12: 401–403.

[37] Lei Y, Zhang X, Su J, Jeong M, Gundry MC, Huang YH, et al. Targeted DNA methylation in vivo using an engineered dCas9-MQ1 fusion protein. Nat Commun 2017; 8: 16026.

[38] Konermann S, Brigham MD, Trevino A, Hsu PD, Heidenreich M, Cong L, et al. Optical control of mammalian endogenous transcription and epigenetic states. Nature 2013; 500: 472–476.

[39] Maeder ML, Angstman JF, Richardson ME, Linder SJ, Cascio VM, Tsai SQ, et al. Targeted DNA demethylation and activation of endogenous genes using programmable TALE-TET1 fusion proteins. Nat Biotechnol 2013; 31: 1137–1142.

[40] McDonald JI, Celik H, Rois LE, Fishberger G, Fowler T, Rees R, et al. Reprogrammable CRISPR/Cas9-based system for inducing site-specific DNA methylation. Biol Open 2016; 5: 866–874.

[41] Saunderson EA, Stepper P, Gomm JJ, Hoa L, Morgan A, Allen MD, et al. Hit-and-run epigenetic editing prevents senescence entry in primary breast cells from healthy donors. Nat Commun 2017; 8: 1450.

[42] Bernstein DL, Le Lay JE, Ruano EG, Kaestner KH. TALE-mediated epigenetic suppression of CDKN2A increases replication in human fibroblasts. J Clin Invest 2015; 125: 1998–2006.

[43] Herman JG, Graff JR, Myöhänen S, Nelkin BD, Baylin SB. Methylation-specific PCR: a novel PCR assay for methylation status of CpG islands. Proc Natl Acad Sci U S A 1996; 93: 9821–9826.

[44] Yu M, Hon GC, Szulwach KE, Song CX, Jin P, Ren B, et al. Tet-assisted bisulfite sequencing of 5-hydroxymethylcytosine. Nat Protoc 2012; 7: 2159–2170.

[45] Deng DJ, Deng GR, Smith MF, Zhou J, Xin HJ, Powell SM, et al. Simultaneous detection of CpG methylation and single nucleotide polymorphism by denaturing high performance liquid chromatography. Nucleic Acids Research 2002; 30: 13E.

[46] Luo DY, Zhang BZ, Lv LB, Xiang SY, Liu YH, Ji JF, et al. Methylation of CpG islands of p16 associated with progression of primary gastric carcinomas. Laboratory Investigation 2006; 86: 591–598.

[47] Zhang B, Xiang S, Yin Y, Gu L, Deng D. C-terminal in Sp1-like artificial zinc-finger proteins plays crucial roles in determining their DNA binding affinity. BMC Biotechnology 2013; 13: 106.

[48] Guo JU, Su Y, Zhong C, Ming GL, Song H. Hydroxylation of 5-methylcytosine by TET1 promotes active DNA demethylation in the adult brain. Cell 2011; 145: 423–434.

[49] Zheng X, Zhou J, Zhang B, Zhang J, Wilson J, Gu L, et al. Critical evaluation of Cbx7 downregulation in primary colon carcinomas and its clinical significance in Chinese patients. BMC cancer 2015; 15: 1172.

